# The Origin and Evolution of Protein Synthesis: A Co-Adaptation Flexible-Rigid Docking Model Based on First-Principles Reasoning

**DOI:** 10.64898/2026.06.11.730790

**Authors:** Dongxu Zhao, Yu Yang, Jian Sun, Jinfeng Zhang, Haotong Duan, Yuxin Tan, Laiyang Liu

**Author notes:** Correspondence and requests for materials should be addressed to D Zhao.

## Abstract

Although the “RNA world” hypothesis suggests that RNA played a crucial role in the origin of life [7], the functional framework of RNA in prebiotic protein synthesis and the mechanisms of genetic code formation during the prebiotic period remain poorly understood. Here, using the prebiotic “primordial soup” as a model, we reconstructed the detailed steps that would yield a protein with a stable ordered amino-acid sequence in the “primordial soup” at the prebiotic period. In the “primordial soup”, a large number of medium- to large-sized biomolecule-like substances—such as RNA-like and protein-like molecules of various sizes and shapes, as well as related polymers like amino-acid-RNA-like etc.—did generate and accumulate. Moreover, protein-like and RNA-like molecules formed even more intricate complexes. These complexes bound free mRNA-like molecules through complementary base pairing. Subsequently, with an extremely low probability, two adjacent amino-acid-RNA-like molecules became bound to this free mRNA-like molecule, and their amino acids underwent a condensation reaction by the complexes, producing peptides and eventually proteins or polypeptides. This free mRNA-like molecule exhibits a certain flexible structure, whereas the super-large complexes formed by protein-like and RNA-like molecules (which possess certain activities) and the amino-acid-RNA molecules exhibit relatively rigid structures. Long-term evolution and mutual selection led to the emergence of proteins with stable amino acid sequences and moderate catalytic activity. In this way, the nucleotide information embedded in such mRNA-like molecules indirectly express through protein synthesis—a process we term the “A Co-Adaptation Flexible-Rigid Docking Model”, where flexible mRNA-like molecules dock onto rigid complexes to enable ordered peptide formation. Finally, we show how trinucleotide codons emerge naturally from the flexible-rigid docking constraints.

---

For more than a century, researchers have never ceased exploring the origin of life and have proposed various hypotheses, such as life originating from a Darwin’s “warm little pond” or the Oparin and Haldane’s “primordial soup.” In addition, the possible formation processes of small biological molecules and biomacromolecules in the prebiotic period have been thoroughly investigated ^[1–6]^. The origin of biomacromolecules such as DNA, RNA, and proteins, as well as the interrelationships among them, is central to exploring or explaining the origin and evolution of life. Although the RNA-world hypothesis suggests that RNA plays a crucial role in the origin of life and protein synthesis ^[7]^, and recent studies have shown that the synthesis of prebiotic peptides also plays an important role in the origin of life [8], researchers still lack a detailed elaboration of the functional framework of RNA in prebiotic protein synthesis, or of how the synthetic pathways of proteins and RNA co-evolved into a complex process with life characteristics during the prebiotic period. Here, we propose a hypothesis for the synthesis process of proteins during the prebiotic period, termed the “co-Adaptation Flexible-Rigid Model,” based on the relationship between the structure and function of biomolecules—particularly focusing on the composition and structural characteristics of enzyme active centers, as well as principles of chemical equilibrium. Furthermore, we deduce the formation of trinucleotide codons based on chemical and biological logic [Fig. 1].

**Figure 1.**
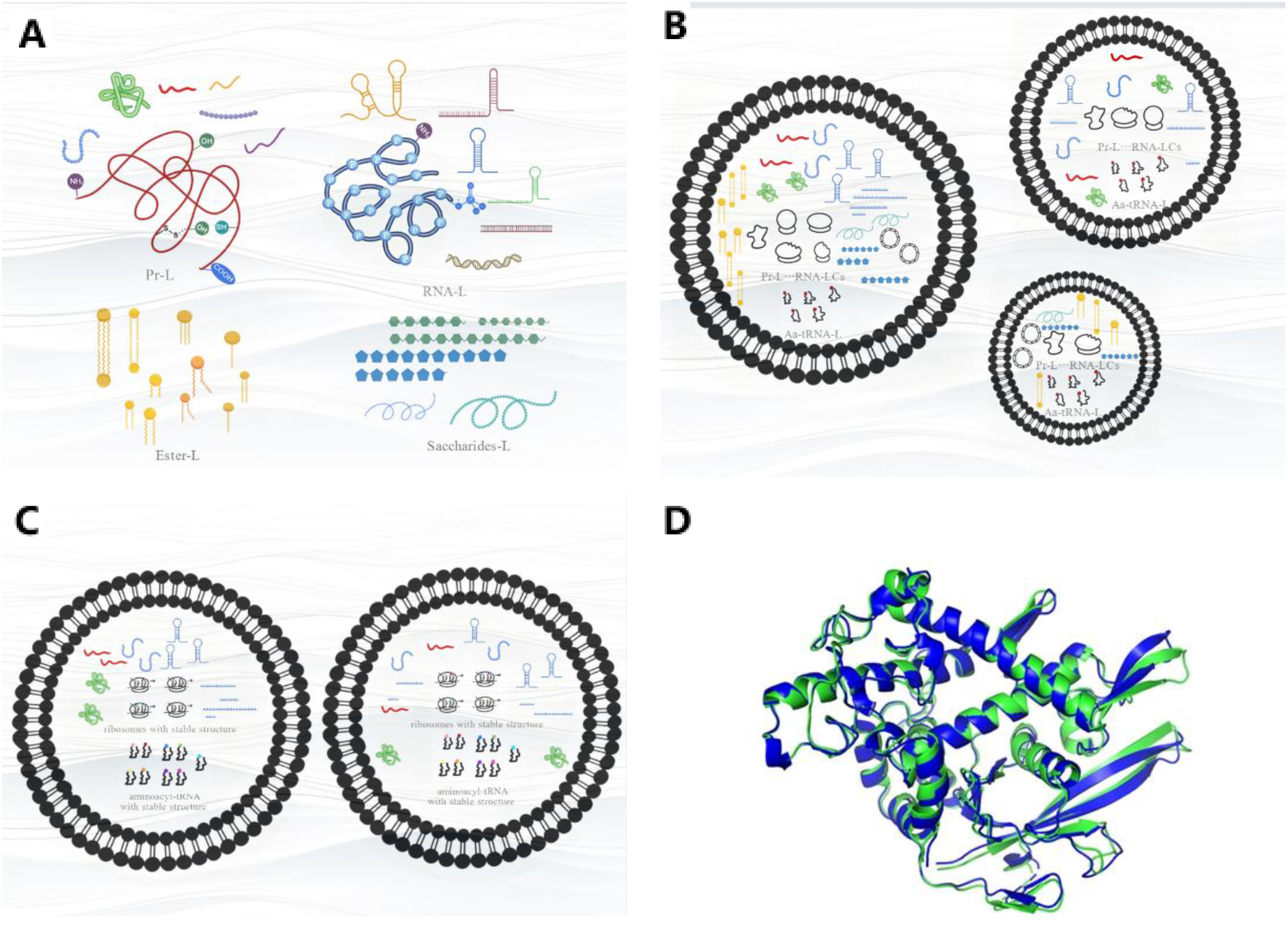
Evolutionary scheme of protein synthesis with stable amino acid sequences in the “primordial soup” during the prebiotic stage. **1A**, the four major types of bio-macromolecules in the “primordial soup”: protein-like, RNA-like, ester-like, and saccharide-like. **1B**, the micro-regions formed by micro-vesicle (The different vesicles encapsulated different bio-macromolecule-like in middle and late prebiotic stage). **1C**, the gradual formation of micro-vesicles with relatively stable compositions of the four major bio-macromolecular components(including stable aminoacyl-tRNA-like, ribosome-like, and mRNA-like molecules) in late prebiotic stage. **1D** proteins from evolution with relatively stable amino acid sequences and structures. Following the formation of the “primordial soup”, various biomolecule-like macromolecules—such as protein-like, RNA-like, ester-like, and saccharide-like substances—were generated through abiotic catalysis. Over time, as the Earth’s temperature decreased, micro-vesicles of various sizes formed in the “primordial soup”, each encapsulating different types of bio-macromolecular analogs. Moreover, the synthesis of protein-like molecules shifted from abiotic catalysis to a primitive, non-canonical form of biological catalysis. Over a long time of evolution, the sizes of these vesicles became increasingly uniform, and meanwhile one new mechanism of the proteins synthesis emerged with stable amino acid sequences and minimal poisonous nature or no harm to the overall system.

## 1 Protein-like or protein-like/RNA-like complexes served as enzyme-like catalytic roles during the prebiotic period

### 1.1 A large quantity of small organic molecule was synthesized via inorganic catalysis or catalysis by organic small molecules at early prebiotic stages

Life on Earth did not exist at the very beginning of the planet’s formation. Instead, it arose through a sequential process: small organic molecules → biomacromolecules → prokaryotic cells. Therefore, from a fundamental logical standpoint, the synthesis of early small organic molecules must have been based on inorganic catalysis and small molecule catalysis. This view is now widely accepted among researchers. Here, we provide a brief overview of this concept.

Based on current geochemical and evolutionary understanding, the Miller-Urey experiment (1953) initially produced various small organic molecules such as amino acids and aldehydes, which subsequently underwent further reactions. Importantly, these findings spurred related research, and subsequent experiments ultimately yielded more complex organic molecules, including monosaccharides, nitrogenous bases, heterocyclic compounds, and nucleotides. As a result, the originally relatively pure water gradually transformed into the so-called “primordial soup” with complex components ^[1–3]^. Therefore, as long as the extreme atmospheric conditions (including chemical composition) and the geodynamic state (e.g., fission) of the Earth did not undergo substantial changes during the transitional period from the late Hadean (4.6–3.8 Ga(gigayears ago)) to the early Archean (3.8–2.5 Ga), these organic synthesis reactions driven by physical factors—such as high temperatures and the Fischer-Tropsch reaction promoted by clay, rocks, etc.—would not cease until the Earth’s environmental conditions fundamentally changed. For example, in 1976, Rohling D.L. found that under high-temperature (65–85°C) and anhydrous conditions, amino acids could polymerize into peptides, and such temperatures were readily achievable in terrestrial environments during the prebiotic period ^[9]^. Clay minerals provided a venue for heterogeneous catalysis, component concentration, and even DNA polymerization, thereby enabling the emergence of early life ^[10–13]^. Studies by Ferris et al. have shown that, in the presence of clay minerals, single-stranded RNA that effectively binds to clay minerals can form longer RNA molecules ^[12,13]^. In addition, mineral nanozymes played an important role during the evolution of organic reactions in the “primordial soup” ^[14–16]^. As a result, the “primordial soup” retained certain concentrations of complex organic molecules, and even various polar molecules could self-assemble into small aggregates under certain conditions. For example, Zhao Y.F. and Benner S.A. focused on the role of phosphorus in the generation of biomolecules ^[17–20]^. Zhao Y.F. et al. synthesized nucleotides by using small molecular N-phosphoryl-α-amino acids and nucleosides as precursors ^[17,18]^, furthermore, the N-phosphorylated amino acids could self-organize into peptides, and even seryl-histidine dipeptides could cleave DNA molecules ^[19]^. Research by Johnston W.K. and other researcher also demonstrates that RNA synthesis from nucleotides can occur without enzymatic catalysis during prebiotic period ^[21]^. Therefore, in the early stages of the prebiotic period, the synthesis of small organic molecules in the “primordial soup” was accomplished through inorganic catalysis and catalysis by small organic molecules.

As mentioned above, a large number of small organic molecules accumulated in the “primordial soup” and the formation of the components that served as structural units of macromolecules predominantly occurred under relatively extreme conditions. As the extreme environmental conditions on Earth diminished, these most primitive synthetic pathways gradually weakened. At that stage, a variety of amino acids, monosaccharides, nitrogenous bases, nucleotides, oligosaccharides, oligopeptides, and oligonucleotides became abundant. During this period, degradation reactions of larger molecules may have occurred to counterbalance the consumption of these small molecules, thereby maintaining their concentration at a certain level.

### 1.2 The transition of bio-macromolecule-like synthesis from catalysis by ordinary inorganic and organic molecules to catalysis by enzyme-like molecules in the “primordial soup”

As previously mentioned, during the mid-stage of the “primordial soup” formation, the various small organic molecules accumulated in the soup would catalyze synthesis and other polymerization reactions among themselves under the relatively extreme physicochemical conditions of that time. This would lead to the generation of oligomers with different degrees and modes of polymerization, such as short peptides, polypeptides, oligonucleotides, oligosaccharides, and polysaccharides of various molecular weights, and would even produce diverse copolymers through the polymerization of different types of molecules. Due to the monotonous and repetitive nature of these reactions, similar polymerization reactions (including limited decomposition reactions) would proceed at a stable rate over a long period until the raw materials were depleted, or until their concentrations remained relatively stable due to chemical equilibrium. To distinguish them from modern proteins and RNA, before the emergence of true prokaryotic cells during the prebiotic period, the involved proteins and RNA are referred to as protein-like (Pr-L) and RNA-like (RNA-L), respectively. Similar notational conventions are used for other biomolecule-like substances throughout this paper. Based on our current understanding of the self-folding properties of biomacromolecules—particularly proteins and RNA—the Pr-L, RNA-L, and Pr-L/RNA-L complexes (Pr-L/RNA-LC) formed through covalent or non-covalent interactions would inevitably adopt specific three-dimensional structures. On the surfaces of these structures, many reactive groups, such as -NH_2_, -COOH, -SH, -OH, imidazole groups, and phosphate groups, would be present. Consequently, some of these structures would exhibit a certain degree of catalytic activity similar to that of biological enzymes—referred to as bio-enzyme-like (Bio-En-L) activity. Although the probability of Bio-En-L occurrence may be low, or even vanishingly low, the abundance of synthetic materials and the sufficiently long time allowed a certain concentration of Bio-En-L to accumulate in the “primordial soup.” This situation may represent what is commonly referred to as a low-probability event. Since these Bio-En-L molecules are not the evolved enzymes we know today, their enzymatic activities or turnover numbers are relatively low. Nevertheless, over the long course of time, life would gradually emerge—a process that simply required patience. In summary, in the early reaction systems rich in small molecular substrates, the reaction process was dominated by polymerization, which generated large quantities of macro-biomolecule-like (MacroB-L) molecules. Furthermore, these early polymerization reactions exhibited characteristics such as substrate randomness, repetitive uniformity, slow reaction rates, and long duration. Reactions promoted by the Bio-En-L components gradually surpassed the ordinary inorganic and organic catalytic pathways, and during the mid-to-late stages of the “primordial soup” or the lower-temperature periods of the Archean, they almost dominated the entire system’s reactions. Regarding the energy consumption of Bio-En-L catalysis, we propose that early Bio-En-L catalysis did consume energy, albeit with high energy consumption or low energy efficiency. Moreover, the energy donors at that time were not necessarily ATP as we know it today; logically, they should have been relatively simple small molecules.

### 1.3 Pr-L is likely to play a dominant role in the biomolecule-like reactions catalyzed by Bio-En-L

Although the organic chemical events we are discussing occurred approximately 4,000 million years ago, because the chemical molecules involved are similar to those in present-day chemistry, we should still apply current fundamental chemical knowledge to understand the chemical reaction mechanisms of that time—including concepts such as the bonding rules of carbon and other elements, as well as the principles of chemical equilibrium. Although Pr-L, RNA-L, and Pr-L/RNA-LC all possess abundant reactive groups, the formation and breaking of bonds during these processes still follow the principles of ordinary nucleophilic and electrophilic reactions. Unlike ordinary inorganic and organic reactions, the Bio-En-L catalytic processes based on Pr-L, RNA-L, and Pr-L/RNA-LC involve active sites that are more complex and random. These active sites do not merely involve a single type of reactive group but rather multiple reactive groups of various kinds. More importantly, during Bio-En-L catalysis, these active sites are also likely to be three-dimensional structures formed by the aforementioned reactive groups.

The discovery of ribozymes provides a new perspective for understanding biocatalytic reactions ^[22,23]^. However, based on our current knowledge of the structures of proteins and RNA, as well as their building blocks, it is necessary to reassess the catalytic role of Pr-L in the middle-to-late stages of the “primordial soup” and its important role in the origin and evolution of life—a role that may even far surpass that of RNA. Pr-L molecules contain active groups such as -NH_2_, -COOH, -SH, -OH, and imidazole groups either in their structures or on their surfaces, and the chemical microenvironments resulting from these groups are relatively mild. Therefore, during the catalytic processes of biomolecule-like macromolecules or Bio-En-L in the “primordial soup,” Pr-L is likely to play a dominant role. In contrast, although RNA-L molecules also possess a certain three-dimensional structure and contain active groups such as -NH_2_, -OH, and phosphate groups that can catalyze some organic reactions, it is precisely the presence of the moderately strong acid—phosphoric acid—that can disrupt the local environment or the microenvironment of the system, thereby destabilizing the stability or even the existence of other biomolecule-like macromolecules. Furthermore, given the simple and repetitive nature of the RNA backbone, its three-dimensional structure is likely unstable (note: the formation of its secondary structure, i.e., local double helices, is somewhat random. For example, a double helix may form between segment A and segment B; similarly, double helices may also form between B and C, A and C, or C and D, which can then randomly give rise to various tertiary structures). Consequently, the Bio-En-L activity of RNA-L varies greatly, making it unsustainable for the formation and maintenance of a stable system.

Based on the above analysis, since we cannot fully replicate the environment and composition of the “primordial soup,” we speculate that during its mid-evolutionary phase (or the Bio-En-L catalytic period), Pr-L, RNA-L, and Pr-L/RNA-LC together played the major catalytic role in the reactions among organic molecules in the “primordial soup” (or the reaction system), making it difficult to distinguish which of the three played the primary or secondary role. However, by no later than the late evolutionary phase, pure Pr-L became dominant and played the primary catalytic role, while pure RNA-L and Pr-L/RNA-LC played secondary roles. Consequently, after long-term evolution, the modern state emerged, as exemplified by ribozyme-mediated splicing of certain RNAs and peptide bond formation facilitated by 23S or 28S rRNA.

Based on the current understanding of protein structure and function, the evolution rule of biomolecules, and other related topics, we argue that rather than viewing the selection of protein-like molecules by life phenomena as the embodiment of life activities as a mere outcome of long-term evolution and selection, it is more accurate to recognize that it was the highly specific structures and high chemical reactivity of proteins that actively promoted the origin and evolution of life.

## 2 Microcompartmentalization during the Mid-Stage of the “Primordial Soup” and the State of Other Biomolecule-Like Molecules Associated with Protein Synthesis

### 2.1 Biomolecule-like macromolecules possessing a certain number of active groups and a certain three-dimensional structure became the “darlings” of the “primordial soup”

During the mid-to-late stages of the “primordial soup,” due to the randomness of reactions, if the macromolecules formed through MacroB-L catalysis were merely inert components, they would play only a minimal role in the evolution of the reaction system, or simply regulate the concentrations of certain small molecular units. In contrast, those biomacromolecules possessing catalytic activity (catalyzing synthesis or decomposition reactions) continually drove the evolution of the reaction system and the continuous change of its components. These active molecules possess a certain three-dimensional structure and contain a certain number of active groups distributed in specific regions. Their level of activity directly affects the stability and evolutionary rate of the entire system. These active biomacromolecules continually induced the production or decomposition of existing macromolecules, while also initiating the synthesis of new macromolecules. This situation is entirely determined by the intrinsic nature of Bio-En-L molecules. Of course, these biomolecule-like macromolecules possessing active groups and a certain three-dimensional structure were also subject to a selection process. Macromolecules that caused minimal or no harm to the system were retained, resulting in the continuous accumulation of active macromolecular populations composed of pure or hybrid polymers, which were themselves also subject to selection. After a long period of mutual selection and adaptation among organic molecules and organic reactions, the biomolecule-like macromolecules, their constituent units, and the derivatives of these units eventually established a coexisting or complementary relationship governed by chemical equilibrium.

In the mid-to-late stages of the “primordial soup,” vesicular structures composed of polar lipid molecules emerged. Within the “primordial soup” and these vesicular structures, the presence, absence, and changing status of the decomposition and polymerization activities of certain Pr-L or RNA-L molecules ultimately determined the trajectory of the system (note: environmental conditions did not undergo extreme changes). When the decomposition and polymerization activities of highly reactive molecules such as Pr-L and RNA-L remained relatively balanced, this not only stabilized the vesicular structures but also maintained the equilibrium of various chemical reactions both inside and outside the vesicles. This process was essentially one of mutual selection between the vesicles and their active components. As a result of long-term evolution, the damage caused by active components to the vesicular membrane gradually decreased. After prolonged evolution, a series of Pr-L, RNA-L, and other similar components—capable of causing minimal disruption to the membrane structure while possessing targeted synthesis and decomposition abilities for their own structures and other components—finally emerged within the complex system of the original vesicular structures, thereby establishing a relatively stable system. This process essentially reflected the biological logic of biocatalysts. We believe that this line of reasoning also provides a new perspective for addressing the long-standing conundrum of whether life originated via a genes-first or metabolism-first approach ^[24–26]^.

In summary, large quantities of medium- to large-sized molecules—such as Pr-L, RNA-L, DNA-like (DNA-L), polysaccharides, and lipid components—were generated and accumulated within the “primordial soup” or vesicle systems during the late prebiotic period. Most of these molecules, including Pr-L and RNA-L, possessed a certain degree of bio-enzyme-like (Bio-En-L) catalytic activity and consequently governed both whether the system evolved and the pace of that evolution.

### 2.2 The vesicles formed by polar lipid molecules endowed the “primordial soup” with characteristics of microcompartmentalization to Some Extent

This section does not discuss the causes, processes, or timing (whether early or late compared to proteins, nucleic acids, and saccharides) of the production of various lipid molecules ^[27,28]^. However, once amphiphilic molecules such as glycerophospholipid-like substances are formed, they inevitably generate relatively stable vesicle structures through hydrophobic interactions in the most common aqueous systems ^[28,29]^. This, to some extent, imparts micro-regionalization characteristics to the originally vast aqueous system, thereby rendering the local microenvironment relatively stable. Of course, this localized micro-regionalization phenomenon also led to a temporary shortage of required raw materials (including energy molecules) within the vesicles. In any case, the formation of lipid vesicles played a profound role in the origin of life. Although researchers have also investigated microspheres composed primarily of proteins or saccharides in the exploration of the origin of life ^[30–32]^, these microspheres could neither easily form structures similar to lipid bilayers nor readily encapsulate the rich components of the “primordial soup”; they tended to form solid structures instead. As a result, they failed to play a central role in the selection process of biomembrane-like components. Current research has made it clear that lipids serve various functions, including energy sources, components of biological membranes, physiologically active substances, surfactants, and solvents. Here, it is necessary to pay tribute to the fundamental property of lipids—their ability to form biological membranes—which provided the most basic structural role in the initiation and evolution of life.

### 2.3 Presence and state of RNA-like molecules in the “primordial soup”

After prolonged chaotic reactions, a vast number and variety of RNA or RNA-L molecules of different sizes, as well as diverse complexes, could form in the “primordial soup”. These molecules included: alcohol-RNA-L, amino acid-RNA-L, and peptide-RNA-L formed via various covalent bonds; RNA-L/RNA-L duplexes (including fully complementary, partially complementary, and other cases); RNA-L/DNA-like hybrid chains (also fully or partially complementary); and RNA-L/Pr-L complexes (including both covalent and non-covalent aggregates). For the larger molecules, unless under extreme physical factors (e.g., high temperature, strong radiation, intense electromagnetic fields) or chemical conditions (e.g., strong acids/bases, high concentrations of metal ions), or a combination of both, their decomposition or formation should rely mainly on the catalytic action of Pr-L and Pr-L/RNA-L (note: not typical biological catalysis). This implies that the probability of population collapse of these molecules remains very low. Aggregation and disaggregation among moderately active molecules occurred from time to time, and only when drastic changes in conditions took place would the entire system undergo noticeable transformation. Important outcomes could include the formation of larger molecular aggregates (analogous to modern biological supramolecular systems) or their complete disappearance, thereby endowing the system with new decomposition characteristics or synthetic properties. Nevertheless, the concentration-dependent chemical equilibrium among these molecules remains the fundamental property of this complex, chaotic system.

### 2.4 Pr-L synthesis during the mid-to-late period of “primordial soup” exhibited significant randomness alongside the evolutionary principles of the system

Proteins, as chain-like molecules composed of amino acids with a repetitive structure, must have been synthesized during the prebiotic era through non-biologically catalyzed polymerization reactions between amino acids or other small molecules containing amino acid residues or scaffolds. Similar to the synthesis of RNA-L described above, the resulting Pr-L was also highly random, monotonous, and repetitive. This was primarily due to the fact that the catalysts present at that time were non-specific — they only took interest in the key/core functional groups (i.e., -COOH and -NH2). With abundant raw materials available, a vast number of protein-like (Pr-L) populations accumulated.

What, then, was the fate of these Pr-L molecules? As analyzed previously, once Pr-L is formed, its relatively stable spatial structure and abundant reactive groups inevitably lead to diversification and rapid changes in the reactions within the system. This makes the system increasingly unstable and even leads to a complete disruption of chemical equilibrium inside the “primordial soup” or within vesicles, thereby causing the system to collapse. Subsequently, the concentration of building blocks that constitute macromolecules reaches a new peak, which then initiates another wave of macromolecular polymer synthesis (including Pr-L, RNA-L, saccharides, lipids, etc.). After countless cycles, a population of moderately active Pr-L eventually emerged and accumulated. Moreover, after long-term selection, new Pr-L and RNA-L finally reached a state of long-term coexistence with new other molecules in the system. Of course, the synthesis pathways of both are still based on the catalytic activity of Pr-L/RNA-L (Note: unless otherwise specified, the term “catalytic activity” here includes the enzyme-like catalytic activity of Pr-L). It must be emphasized again that the synthesis of Pr-L is not based on the genetic logic of specific sequences of RNA-L — in other words, the synthesis of Pr-L has no genetic characteristics.

## 3 Synthesis of Protein-like Molecules with Specific Amino Acid Sequences in the “Primordial Soup”

The previous text analyzed the random synthesis and decomposition of RNA-L and Pr-L. The following section focuses on the framework and rationale for the synthesis of Pr-L with a specific amino acid sequence during the late stages of the “primordial soup” — namely, the biological and chemical logic of codon formation.

### 3.1 The possibility of numerous RNA-L, amino acid-RNA, and other medium to large molecules being involved in peptide/protein-like synthesis

The formation of supermolecules based on RNA-L structures in the “primordial soup” was inevitable. Under ordinary conditions, base complementarity would inevitably cause numerous pure RNA-L molecules and other components containing RNA-L motifs to form aggregates of various sizes, which might be pure or mixed. Moreover, these mixed aggregates likely accounted for a large proportion of all aggregates formed. In the absence of intense mechanical disturbances, high temperatures, and other factors, only the relatively simple chemical environments (i.e., non-strong acids, non-strong bases, high ionic strength, etc.) could drive the formation of super-large and disordered aggregate. It is now well understood that, with the aid of various weak interactions including base complementarity, biomolecules can easily form aggregates of different sizes. For example, in current processes for preparing bio-based nanoparticles or microspheres, techniques such as magnetic stirring, ultrasonic treatment, microporous extrusion, and various high-speed emulsification operations are required to disperse the aggregates and achieve the desired size range ^[33-35]^. Consequently, when characterizing aggregate stability, it is still necessary to observe whether precipitation (i.e., re-aggregation of micro- and nanoparticles) occurs over a certain period of time.

Similar to the rRNA structures known today, the resulting aggregates, predominantly composed of RNA-L, exhibit a certain spatial configuration. Although this structure is generally a relatively dense entity, spaces, cavities, or even gaps do exist within or on its surface. Moreover, because nucleotides are larger and more complex in structure than amino acids, the compactness (note: not density) of three-dimensional entities mainly composed of RNA-L is lower than that of proteins. Due to the prevalence of three-dimensional structures among the aggregates described above, as well as the presence of numerous active groups (e.g., -NH2, -COOH, -SH, -OH, imidazole groups, etc.) and inert groups (e.g., various aliphatic chains of different lengths, benzene rings, etc.), these aggregates can also exhibit enzyme-like catalytic characteristics. During or after the formation of these aggregates, active groups located at specific sites/positions are likely to trigger certain group transfer reactions, thereby disturbing the existing structure and reconstructing the local structure. If two amino acid residues connected to RNA happen to be present at that specific site, a transfer of amino acid residues may occur, thereby forming the so-called peptide bond. Of course, due to the randomness of the amino acid-RNA molecules involved in the aggregates, or the random diffusion of the amino acid moieties of two amino acid-RNAs into specific cavities, the formation of peptide bonds is highly random. Moreover, such a super-large aggregate can also facilitate the synthesis of other molecules or derivatives.

This type of super-large aggregate differs from the enzyme-like catalysis of the numerous Pr-L…RNA-L complexes in the aforementioned composite system. The latter primarily catalyze reactions among relatively small molecules, whereas the former can trigger reactions involving molecules that are temporarily fixed to the aggregate (via base complementarity). This work of super-large aggregate differs from the previously mentioned enzyme-like catalysis of numerous Pr-L/RNA-LCs. The latter primarily catalyze reaction among relative small molecules, whereas the former only can trigger reactions involving molecules that are temporarily fixed to the aggregate (via base complementarity). Although no research has confirmed the existence of the super-large aggregates, the known complexity of the spliceosome involved in intron splicing offers some perspective. The spliceosome contains more than 140 kinds of proteins and 5 kinds of snRNAs (totaling about 10,000 nucleotides). Is every single component useful? The answer is clearly no. We believe that the presence of most of these components in the spliceosome merely has a minimally harmful or negligible effect on cellular metabolism. This brings to mind the modern ribosome, which also contains numerous protein components, most of which (primarily proteins) are merely evolutionarily retained redundancies.

### 3.2 Possible mechanisms for the formation of polypeptides or Pr-L with a given stable amino acid sequence in the “primordial soup”

Stable synthesis of peptides/Pr-L implies the consistent reproduction of amino acid sequences. Based on current knowledge, Pr-L and nucleic acids play essential roles in life processes, especially in genetic processes. However, in the mid-stage of “primordial soup” or stage of vesicles, when it comes to their contribution to the structural stability of the system, their roles are less important than those of membrane lipids. As mentioned earlier, in the mid-stage of “primordial soup” system (including vesicle contents; this applies throughout), a population of components with relatively moderate activity, such as Pr-L and RNA-L, eventually came to persist. Over time, the existing moderately active Pr-L would gradually decompose due to environmental changes or the instability of their own covalent structures. So, in such a complex reaction system, how can these moderately active Pr-L be continuously regenerated?

Although we now have a relatively clear understanding of the mechanisms of protein synthesis within cells, particularly how mRNA and tRNA participate in this process, we still know very little about how the random polymerization of amino acids during the prebiotic era evolved into the state observed in cellular systems. Here, based on the catalytic activity of the aforementioned aggregates mainly formed primarily with RNA-L as the polymerization scaffold, we propose a possible mechanism for the synthesis of proteins with stable amino acid sequences — the Flexible-Rigid co-Adaptation hypothesis (also referred to as the Flexible-Rigid co-Adaptation Model) (Fig. 2). This mechanism is based on at least two underlying principles:

**Figure 2.**
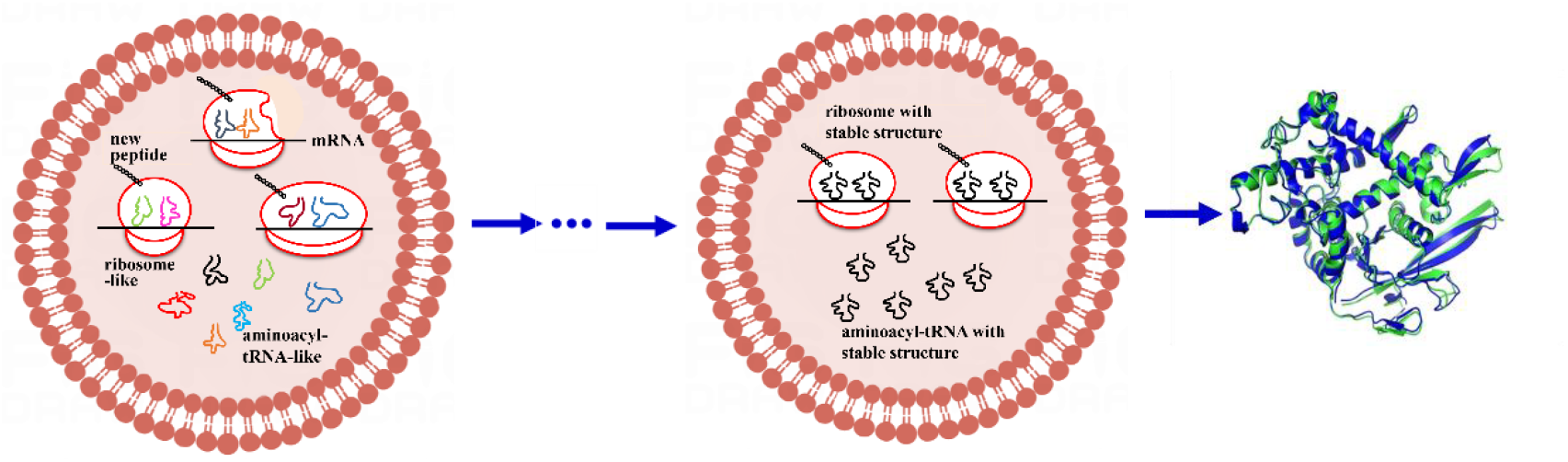
Schematic diagram of the evolutionary pathway for the protein synthesis with stable amino acid sequence based on super-large Pr-L/RNA-L complexes in the prebiotic stage(Flexible-Rigid co-Adaptation Model)

Within the vesicle-like structures, various forms of super-large Pr-L/RNA-L complexes (i.e., ribosome-like) and mRNA-like molecules, aminoacyl-tRNA-like molecules are assembled into diverse apparatuses to synthesize corresponding peptide chains at a relatively slow rate(Left). Over long periods of evolution, due to the relative stability of the linear or flexible structures of mRNA-like molecules, the ribosome-like and aminoacyl-tRNA-like molecules that were somewhat complementary to them were selected. Moreover, the latter two underwent a more active mutual selection, aiming for the ribosome-like molecules to fully accommodate the aminoacyl-tRNA-like molecules. This process eventually evolved into the current state: the structures of ribosomes and aminoacyl-tRNAs are relatively stable, and their spatial configurations become fully complementary upon binding with mRNA (Middle).

### 1) A fundamental principle underlying sequence-stable peptide/protein synthesis is the requirement for a template

Unlike homopolysaccharides, which consist of uniform monosaccharide units, the numerous amino acids, upon entering a specific assembly workshop (synthesis site), must have specifically differentiated spaces, sites, or a template to recognize and position different amino acids. In other words, the template must possess the ability to select among different amino acids, and moreover, it must be sufficiently large or long — a point that is very important.

### 2) The second principle for sequence-stable peptide/protein synthesis is that amino acids should be rendered inert, i.e., tagged, before participating in reactions

Amino acids are small molecules and thus have very high diffusivity in aqueous systems, which makes it difficult for two amino acid molecules to collide and become temporarily fixed at specific sites within the ultra-large Pr-L/RNA-LC.

Therefore, other molecules — such as shorter RNA chains, oligosaccharide chains, or long-chain lipid molecules — can render amino acids inert. Among these, the use of shorter RNA-L is most advantageous: it not only increases the macroscopic size of the amino acid but also enables selective temporary fixation through base-pairing with the RNA component of the ultra-large Pr-L…RNA-LC, thereby increasing the contact time between two amino acids and facilitating the reaction. It is known that at a temperature of around 297 K, the diffusion coefficient D of amino acids in the aqueous phase is approximately 1.0 × 10^−5^ cm^2^/s, while the diffusion coefficients of proteins and RNA are slightly less than 1.0 × 10^−6^ cm^2^/s, and for large molecules, the value is only about 10^−7^ cm^2^/s ^[36-38]^. Therefore, the mobility of amino acid-RNA-L (also called aminoacyl-tRNA-like) molecules is significantly lower than that of free amino acids. Consequently, in the mixed system, the probability that two amino acid-RNA molecules simultaneously bind to the RNA-based aggregates increases significantly. Interestingly, this inertization process also endows a given amino acid with a specific tag — namely, the codon (which will be elaborated later).

Based on the above analysis of the two basic logics required for protein-like synthesis, a possible mechanism for reproducible protein synthesis is as follows: A long RNA in the system (referred to as “target mRNA-like”) temporarily binds, through limited base interactions, to a super-macromolecular Pr-L/RNA-LC (a biological catalyst-like molecule), forming a binary complex—super-large Pr-L/RNA-LC (ribosome-like)/mRNA-like template. Meanwhile, two aminoacyl tRNA-like molecules in the system also happen to bind, through base complementarity, to the same target mRNA-like. Consequently, the attached amino acids are brought into close proximity in space and remain so for an extended period, thereby forming a transient quaternary complex. In this quaternary complex, the super-macromolecular Pr-L/RNA-LC acts as a catalyst, enabling the first template-dependent synthesis of Pr-L — i.e., the formation of peptide bonds using the “target mRNA-like” as a template — at the proto-life stage. (Note: This reaction was not the first bio-synthesis-based mode.). The detailed explanation is as follows:

### 1) Randomness and diversity of the formed quaternary complexes

In the middle and late stages of the “primordial soup”, the small aminoacyl tRNA-like molecules did not bind only to the target mRNA-like when interacting with the binary complexes; they also had a significant probability of binding to the RNA-L component of the ultra-large Pr-L/RNA-LC. In such cases, however, no new peptide bond would form. In other words, the quaternary complex exhibited high compositional and structural diversity. Two similarly sized aminoacyl tRNA-like molecules may bind to the same mRNA-like, with their attached amino acids positioned close to each other and with catalytically active groups nearby. Although the probability of such a specific configuration is very low, it is precisely this low-probability event — combined with its long duration — that enables the frequent occurrence of an otherwise rare event, thereby leading to the stable synthesis of proteins with specific amino acid sequences. Researchers randomly synthesized an enormous variety of RNA molecules (exceeding 10^14^ different types) using combinatorial chemistry techniques. From this pool, 16 kinds of RNA molecules capable of binding ATP were obtained by affinity chromatography. When these RNAs bound to ATP, they showed both high specificity and high affinity (with dissociation equilibrium concentrations below 50 µmol/L, far lower than typical intracellular ATP levels of 2–10 mmol/L) ^[39]^, thereby empirically supporting the above hypothesis. This result also aptly demonstrates that, in the early random reactions closely associated with the origin of life, certain biomacromolecules with specific structures and functions can always be produced. Although the probability of their emergence is extremely low, it is precisely the result of natural selection.

### 2) The initially formed binary complex (the super-large Pr-L/RNA-LC (ribosome-like)/mRNA-like template) exerts a screening effect on the bound aminoacyl-tRNA-like molecules according to their size and shape

The initially formed binary complex is only one of many complexes in the “primordial soup”. Only when the two entering aminoacyl-tRNA-like molecules have very similar sizes or shapes can their attached amino acids approach each other within the limited physical space. In other words, due to its relatively fixed structure, this catalytically active super-large Pr-L/RNA-LC exhibits a strong screening effect on the many aminoacyl-tRNA-like molecules in the system according to their size and shape.

In the mid-stage of the “primordial soup”, among the population of amino acid-RNA-like molecules, the single amino acid-single nucleotide pairing would theoretically have been most suitable — most easily to put the amino acid-RNA-like molecules to the target site. This scenario would have required the presence of nucleotides equal in number to, or slightly exceeding, the number of amino acid types. However, although single amino acid-dinucleotide or single amino acid-trinucleotide compositions could accommodate different amino acids through various nucleotide combinations, the rigidity of short chains (e.g., Phe-UUU, Phe-UUC, etc.) or their resistance to conformational change made it difficult for the attached amino acids (which had already been directed) to come into close physical proximity (i.e., enforced proximity). Consequently, the transition from intermolecular reactions to intramolecular reactions was difficult to achieve. How could this dilemma be resolved? In the process of molecular evolution, the strategy of using a relatively large RNA structure to temporarily fix amino acids was adopted — namely, the L-shaped three-dimensional structure of present-day aminoacyl-tRNA.

The evolutionary logic governing the shape of aminoacyl-tRNA-like molecules is as follows: Since two aminoacyl-tRNA-like molecules temporarily coagulate inside the ultra-large Pr-L/RNA-LC via diffusion, smaller folded structures would have been more advantageous from a basic logical perspective. Ideally, it should be rod-shaped, with a protruding single-stranded segment to facilitate amino acid attachment, and should allow significant twisting so that the two attached amino acids can be brought closer together. Although modern tRNA does not have a typical rod shape, its compact L-shaped structure compresses its volume to the extreme, and its appearance can be regarded as rod-like.

Because the active sites and structure of the catalytic Bio-En-L (the super-large Pr-L/RNA-LC) are relatively fixed, it must both maintain moderate interaction with the mRNA-like template and position the two attached amino acids as close together as possible in space. Long-term selection resulted in the gradual enrichment of components resembling modern tRNA, ultimately preserving a class of molecules with specific structures similar to those of modern tRNA (though not necessarily the familiar cloverleaf shape). Other types of aminoacyl-tRNA-like molecules whose structures were not clearly recognized by the super-large Pr-L/RNA-LC gradually accumulated, which to some extent fed back negatively into their synthesis, thereby leading to their gradual elimination. At the same time, this process also meant that the super-large Pr-L/RNA-LC itself was subject to selection. The mutual selection process among the super-large Pr-L/RNA-LC, aminoacyl-tRNA-like molecules, and mRNA-like molecules was essentially the evolutionary process during the prebiotic era. As Lewontin R.C. (1978)once said, “Evolution is best viewed as a history of organisms finding devious routes around constraints” ^[40]^. The reason that early repeated organic reactions were able to give rise to life phenomena is essentially that the system was merely selecting or finding a way to sustain itself over extended periods.

### 3) The conformation lability or structural flexibilituy of the ultra-large Pr-L/RNA-LC

The ultra-large Pr-L/RNA-LC was also subject to a self-elimination process during its long-term catalysis of new peptide bond formation. Beyond possessing enzyme-like catalytic activity, it also had a certain degree of mobility — or, in other words, significant structural flexibility or lability — in its local regions. Peptide bond formation inevitably causes changes in spatially adjacent local groups. The ultra-large Pr-L/RNA-LC cannot remain completely static; otherwise, it would form a non-functional quaternary complex, and subsequent peptide bond formation would be halted. Overly rigid ultra-large Pr-L…RNA-LC molecules were gradually eliminated. Peptide bond formation inevitably involves the transfer of amino acid residues from two aminoacyl tRNAs, as well as the storage and export of the growing product. This process undoubtedly induces significant local structural adjustments or displacements within the ultra-large Pr-L/RNA-LC. Obviously, for a molecule held together entirely by covalent bonds, it is difficult to undergo such large spatial displacements and to accommodate the ‘huge product’. Therefore, transient complexes formed by multiple components, owing to their non-rigidity and mobility, reduced the resistance during amino acid residue transfer. The long-term outcome of this evolutionary process is the familiar composition of the modern ribosome, consisting of large and small subunits.

### 4) Mutual selection and elimination between ultra-large Pr-L/RNA-LC and mRNA-like molecules

In the mutual selection process between the ultra-large Pr-L/RNA-LC and free mRNA-like molecules, a short conserved sequence capable of base pairing became essential for both parties. For mRNA-like molecules, this short sequence is what is now called the SD-like sequence. RNA-L molecules with SD-like sequences were more likely to be preserved and to have their encoded information reproduced in protein form. Similarly, Pr-L…RNA-LC molecules that contained sequences complementary to the SD-like sequence were also retained. Since the interaction between the two molecules is temporary, the complementary base pairs formed should not be too long; ideally, they should not easily form a double helix, and the pairing region should preferably be shorter than 5-6 bp. We suggest that the reason the current SD sequence serves as a conserved sequence for both parties is likely related to the need to maintain a certain degree of stability between the mRNA-like molecule and the Pr-L/RNA-LC. Thus, after the mRNA-like molecule binds stably to the Pr-L/RNA-LC, this stable complex provides sufficient time for the sequential binding of the two aminoacyl-tRNA molecules to the mRNA-like molecule. Moreover, the ‘…AGGAGG…’ arrangement has certain distinctive features. In fact, other arrangements — such as ‘…TGGTGG…’, ‘…GGAGGA…’, ‘…TCCTCC…’, or ‘…GGTGGT…’ — might also be acceptable, provided that the corresponding sequence in the Pr-L/RNA-LC is adjusted accordingly. After the formation of the first peptide bond, the super-large Pr-L…RNA-LC must dissociate from the conserved sequence of the mRNA-like molecule to facilitate subsequent peptide bond formation. This dissociation requires energy, which is currently generally supplied by GTP hydrolysis.

The energy required for the dissociation of the super-large Pr-L/RNA-LC from the SD-like sequence of the mRNA-like molecule is estimated as follows: The strength of hydrogen bonds in base pairs is greatly affected by molecular structure and environmental conditions, generally ranging from 4 to 21 kJ/mol. Considering the imperfect formation of hydrogen bonds in base pairs during the late prebiotic period (note: at this stage, the hydrogen bonds formed between two larger molecules may not have been perfectly spatially matched), we take the hydrogen bond energy at a low value of 4 kJ/mol, and each base pair is assumed to have an average of 2.5 hydrogen bonds. Therefore, the total hydrogen bond energy per base pair ≈ 4 (kJ/mol) × 2.5 = 10 kJ/mol. Since the bond energy of a single high-energy phosphate bond is 30.54 kJ/mol, and 30.54 kJ/mol / 10 kJ/mol ≈ 3, the hydrolysis of one high-energy phosphate bond from GTP can dissociate approximately 3 base pairs. Moreover, since this energy consumption occurs only at the initiation of protein synthesis, consuming even two high-energy phosphate bonds is quite necessary and, in fact, very economical for a single round of protein synthesis. The current SD sequence is typically 4–9 nucleotides long. Considering the pulling effect of the large subunit’s movement on the small subunit, the energy required to separate the SD sequence from the small subunit should not exceed two high-energy phosphate bonds.

### 5) The Mechanism of Codon Formation

How many base pairs are involved in the interaction between an aminoacyl tRNA-like molecule and a random mRNA-like molecule? This question essentially addresses the number of base pairs in such interactions, which fundamentally determines the mechanism of codon formation. The purpose of the interaction between an aminoacyl tRNA-like molecule and a free mRNA-like molecule is to temporarily bring two amino acids into close physical proximity. This process involves at least two key issues: first, the number of base pairs required for the interaction; and second, the size of the tRNA-like molecule (to be analyzed in detail below). Regarding the first issue, from a fundamental logical perspective, since aminoacyl tRNA-like molecules temporarily and randomly fix amino acids at specific sites and must be able to leave easily, it is appropriate to have 1–4 interacting base pairs. In the limited space within the ultra-large Pr-L/RNA-LC, the mRNA-like molecule can more easily maintain a quasi-linear structure for pairing with each aminoacyl tRNA-like molecule, and after base pairing, the formation of stable double helices must be prevented. Regardless of whether the dissociation of the aminoacyl tRNA-like molecule is a passive or active process, disrupting the formed base pairs is a process that requires energy, either directly or indirectly. The energy consumption of this process should be slightly greater than an integer multiple of the binding energy of a single base pair (10.0 kJ/mol), i.e., greater than 10.0, 20.0, 30.0, or 40.0 kJ/mol. Based on our current understanding of bioenergetics of metabolism, this energy consumption should be less than the energy contained in one ATP (the bond energy of a single high-energy phosphate bond is 30.54 kJ/mol). Therefore, from a bioenergetic perspective, the energy required for an aminoacyl-tRNA-like molecule to dissociate from the mRNA-like molecule or from the ultra-large Pr-L/RNA-LC should be close to 30.54 kJ/mol. According to our current understanding of codons, the binding energy of the third base pair of a codon differs from that of the first two base pairs. If we estimate this binding energy as 75% of the normal bond energy, then the binding energy between a codon and its anticodon is 10.0 kJ/mol × 2.75 =27.5 kJ/mol, meaning that the energy from one ATP molecule is sufficient to dissociate one codon. In fact, as is well known, after the formation of a new peptide bond, the introduction of a new aminoacyl-tRNA consumes one GTP (equivalent to one ATP), which is entirely sufficient to dissociate one codon. That is to say, from an energy balance perspective, the system is self-consistent, with only a small amount of energy redundancy. For current cellular metabolism, this can already be considered quite intelligent.

The analysis of amino acid orientation by the aminoacyl-tRNA-like molecule based on its quasi-linear complementarity with the mRNA-like molecule is as follows: Because mRNA-like molecules are quasi-linear and connected by single phosphodiester bonds, they exhibit considerable flexibility. When an aminoacyl tRNA-like molecule (analyzed previously) with a three-dimensional structure binds to it via only a single base pair, the spatial position of the tRNA-like molecule or the spatial orientation of the attached amino acid, becomes uncertain. Consequently, 2–3 base pairs are necessary to properly orient the amino acid (Fig. 3). Moreover, due to the relatively confined space within the ultra-large Pr-L…RNA-LC, not only is it difficult for the mRNA-like molecule to maintain a linear structure, but it is also difficult for three consecutive nucleotides in the three-dimensional tRNA-like molecule to maintain a linear arrangement (except at the termini). This makes the binding between the two local linear structures somewhat difficult. Thus, according to basic geometric principles, two points can easily define a straight line, whereas three points are less likely to be collinear but can define a relatively stable plane. Thus, three base pairs provide the necessary spatial orientation for the two amino acids.

**Figure 3.**
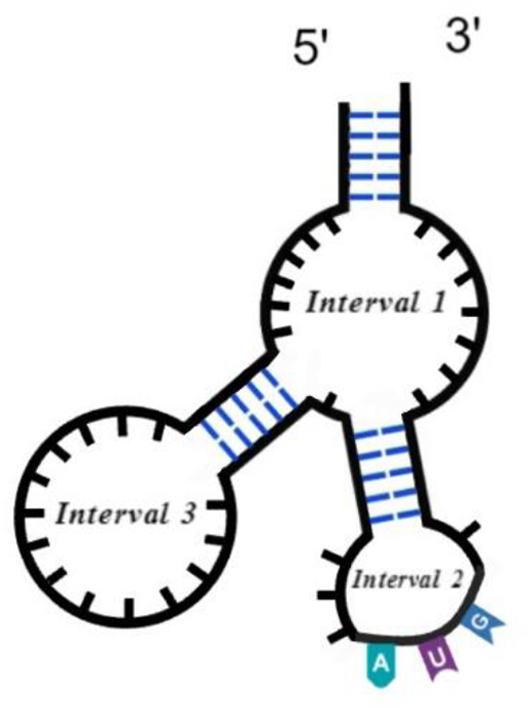
Schematic diagram of the theoretical size of aminoacyl-tRNA-like molecules. The tRNA-like molecule contains three double-helical segments, each composed of 5 base pairs, separated by single-stranded regions of 7 nucleotides. The upper part, where the amino acid is attached, contains several unpaired nucleotides, while the lower part is the region that pairs with mRNA-like molecules.

One possible mechanism is as follows: The tRNA-like chain of the aminoacyl-tRNA-like molecule should have a three-dimensional structure with considerable flexibility or torsional capability. At the same time, after complementary base pairing has occurred, there should be minimal bending between the first base pair of the first aminoacyl-tRNA-like molecule and the third base pair (i.e., the terminal base pair) of the second aminoacyl-tRNA-like molecule. Coupled with the twisting of the tRNA-like three-dimensional structure and the torsion of the nucleotides directly attached to the amino acid, this mechanism brings the two amino acids as close together as possible. This analysis has considered the number of base pairings between aminoacyl-tRNA-like and mRNA-like molecules from a geometric perspective. Next, we will address the issue from the perspective of energy consumption.

The entry of the aminoacyl-tRNA-like molecule into the complex is a free process driven by diffusion and does not consume energy. Once inside, it must be transiently stabilized within the complex through base-pairing with the mRNA-like molecule — a process that we consider to be energy-consuming. As is well known, during protein synthesis, the entry of a non-initiator aminoacyl-tRNA into the ribosome consumes one GTP (equivalent to one ATP), which is likely related to this energy requirement. Since the hydrogen bond energy of each base pair is approximately 10 kJ/mol, the hydrolysis of one high-energy phosphate bond can stably bind up to three base pairs. From this, the current codon structure (three nucleotides per codon) can be deduced. Moreover, because it is difficult for the aminoacyl-tRNA to maintain a local linear structure, the third base pair formed after binding to the mRNA may be in a less tightly associated state. In other words, the third base of the participating mRNA-like molecule cannot easily maintain its spatial orientation, whereas the third base of the aminoacyl-tRNA (which is now the first base of the anticodon) has a defined spatial orientation. As a result, the binding energy of the third base pair is weaker than that of the first two, exhibiting a certain degree of non-stringency. As is well known, the wobble of the third base pair in codon-anticodon pairing may be related to this phenomenon. As for the purely mathematical explanation based on 20 amino acids and 4 nucleotides — namely, that three consecutive nucleotide combinations (4^3^ = 64) fully satisfy the requirements for 20 amino acids — this explanation appears plausible on the surface but is actually far-fetched.

Coincidentally, and with a hint of predestination, three consecutive nucleotides can generate 64 distinct groups — i.e., they can accommodate up to 64 different amino acids.

Interestingly, because the number of amino acid types preserved in proteins through long-term evolution is smaller than what the genetic code could theoretically accommodate, the well-known phenomenon of codon degeneracy has emerged (not elaborated here).

### 6) Theoretical estimation of the minimum number of nucleotides required for a stable three-dimensional aminoacyl-tRNA-like molecule

Previously, we made a general prediction regarding the size and structural principles of aminoacyl-tRNA-like molecules. But what, then, is the appropriate size for such a molecule?

Here, we draw on the concept of protein super-secondary structure. The major forms of protein super-secondary structure include αα, βαβ, and βββ — meaning that two or three typical secondary structures are required to form a minimal and stable higher-order structure. Therefore, we hypothesize that the three-dimensional RNA entity of the aminoacyl-tRNA-like molecule is composed of at least three double-helical segments.

Assuming that local double-helical regions account for approximately 45% of a typical RNA molecule, with each helix containing only 5 base pairs, a spacer of 7 nucleotides between helical segments, the interaction site with the free mRNA-like molecule located at one end of the three-dimensional structure, and the amino acid-binding helix at the opposite end, the required number of nucleotides is approximately (starting from the 5′ end):

5 nt (Helix-1) + 7 nt (Spacer-1) + 5 nt (Helix-2) + 7 nt (Spacer-2) + 5 nt (Helix-3) + 7 nt (Spacer-3, unpaired, interacting with the free target mRNA-like) + 5 nt (Helix-3) + 7 nt (Spacer-2) + 5 nt (Helix-2) + 7 nt (Spacer-1) + 5 nt (Helix-1) + 3 nt (amino acid-binding end) + 7 nt (10% redundancy) = 75 nt (Fig. 3).

This estimate is remarkably close to the size of modern tRNAs, which generally range from 70 to 90 nucleotides.

In fact, stable protein synthesis reflects its intrinsic biological logic [41]. Since the sequence of amino acids and the peptide bonds (or peptide planes) are fundamental characteristics of proteins, this naturally leads us to consider further that the synthesis of a specific protein could be accomplished by a giant device equipped with a long groove or channel — which can be called a “long workshop [with a predetermined template]” — capable of recognizing and accommodating different amino acids. Obviously, nature would not evolve countless such protein-synthesis devices to accomplish protein synthesis; that is, a long workshop with a certain degree of rigidity or hardness should be implausible. Could a stable double-helical DNA molecule serve as a long template for housing amino acids? Due to the monotonous repetitiveness of DNA structure, even if it could trigger condensation reactions through transient fixation of amino acids with the assistance of other biological macromolecular aggregates, the limited variety of base pairs in DNA would be insufficient to handle the many types of proteinaceous amino acids — far more than the familiar 20 or so.

The resulting proteins would either be repeats of limited amino acid units or completely disordered polymeric amino acids, thus failing to solve the key problem of sequence stability. By the same logic, the combination of a movable small workshop and a long rigid template — the so-called ‘small workshop model’ — is also infeasible. Therefore, due to the necessity of a ‘long template’, perhaps the only remaining option in the evolution of protein synthesis is the pairing of a ‘rigid’ synthesis device (a small workshop) with a ‘flexible’ template. In essence, this establishes the relationship between flexible RNA and a rigid protein synthesis device.

The origin and evolution of the genetic code is a subject of considerable interest. As early as 1965, Woese C.R. proposed a hypothesis, known as the stereochemical hypothesis, suggesting that the basis for the emergence of the genetic code may lie in a potential chemical matching between amino acids and trinucleotide codons, or that some amino acids have a selective chemical binding affinity with their corresponding codons ^[42]^. Conversely, F.H.C. Crick (1968) held another view. He proposed that codons originated from a ‘frozen accident’, suggesting that there is no necessary chemical link between codons and their corresponding amino acids; rather, the association is random ^[43]^.

To this day, the origin of codons remains unresolved. Earlier, we analyzed the origin of codons in some detail, but did not address the relationship between codon composition and the corresponding amino acids — or, more specifically, why the codons UUU and UUC both represent phenylalanine (Phe) rather than the structurally similar tyrosine (Tyr). Based on the above analysis, we propose that the correspondence between early codons and amino acids was indeed random. However, as mutual selection occurred among amino acids, aminoacyl-tRNA-like molecules, and ribosome-like complexes, the number of amino acid types participating in protein synthesis gradually decreased, giving rise to characteristics such as codon degeneracy and wobble. Meanwhile, we also suggest that codon wobble is, to some extent, a “tolerant” strategy adopted to accommodate both the linearity and flexibility of mRNA and the local regions of aminoacyl-tRNA-like molecules — which possess a certain three-dimensional structure but can hardly maintain a long linear arrangement for pairing.

## 4 Conclusion and Outlook

After a long period of selection, a mutually compatible state among the ultra-large Pr-L/RNA-LC (ribosome-like), the mRNA-like template, and two aminoacyl-tRNA-like molecules (aminoacyl-tRNA1 and aminoacyl-tRNA2) finally emerged in the “primordial soup” or vesicles. As evolution proceeded, the variety of ultra-large Pr-L/RNA-LC types gradually decreased, as did the variety of mRNA-like templates (later, with the emergence of cellular forms, proteins determined by the system and harmless to the system gradually emerged and were preserved). Similarly, the types of aminoacyl-tRNA-like molecules also decreased to match the declining ultra-large Pr-L/RNA-LC. Specific proteins were fairly well regenerated based on the template RNA, i.e., the mRNA-like nucleotide sequence, or in other words, the information embedded in the specific nucleotide sequence of the template RNA was indirectly reproduced. The ultra-large Pr-L/RNA-LC and aminoacyl-tRNA-like molecules exhibit a certain degree of rigidity, whereas the target mRNA-like molecule (template RNA), existing in a linear form, displays a certain degree of flexibility. Through the bases of the aminoacyl-tRNA-like molecules, a movable docking between the rigid and flexible structures is achieved. This is the “Flexible-Rigid co-Adaptation model” that allows a specific protein to be reproduced within vesicles.

Although researchers have fortunately obtained very limited fossil samples from the earliest stages of life ^[44-47]^, which have pushed the earliest emergence of life back from 3.5 Ga to 3.95 Ga, we are still at a speculative stage regarding the formation and evolution of early biological macromolecules. We cannot yet fully replicate in the laboratory or test tube the actual reactions that occurred during the darkest period of life’s origin. Thus, the synthesis of protein-like molecules or proteins during the prebiotic period can only be hypothesized and experimentally investigated based on our current knowledge of protein synthesis.

Thanks to combinatorial chemistry, we believe that by using independently randomly synthesized peptide libraries, independently randomly synthesized oligonucleotide libraries, and mixtures of the two as potential Bio-En-L (Note: under certain conditions, the given system may possess Bio-En-L functionality), and by using individual nucleotides, individual amino acids, or mixtures thereof as substrates, we can alter experimental conditions such as temperature, pH, and specific metal ions. Through repeated experiments, we hope to obtain biomolecular supramolecular assemblies or specialized reaction systems with specific catalytic functions. This would allow us to gain insight into the types of reactions and corresponding products that may have existed during early evolution, thereby complementing theoretical conjectures and simulations.

Protein synthesis in the prebiotic period is a subject that is exceedingly remote, profound, complex, and precise. Moreover, the associated early chemical reactions were random or contingent. It was only after long-term evolution and selection that the interactions among biomolecules gradually became restricted and directed.

## Author Contributions

D.ZHAO conceived and designed the study, and wrote the manuscript. Y YANG, J SUN and J ZHANG gave the valuable suggestion and refined this manuscript. H DUAN, Y TAN and L LIU collected the relative articles and prepared the figures.

